# Conditional GWAS analysis identifies putative disorder-specific SNPs for psychiatric disorders

**DOI:** 10.1101/592899

**Authors:** Enda M Byrne, Zhihong Zhu, Ting Qi, Nathan G Skene, Julien Bryois, Antonio F Pardinas, Eli Stahl, Jordan W Smoller, Marcella Rietschel, Bipolar Working Group of the Psychiatric Genomics Consortium, Major Depressive Disorder Working Group of the Psychiatric Genomics Consortium, Michael J Owen, James T.R. Walters, Michael C O’Donovan, John G McGrath, Jens Hjerling-Leffler, Patrick F Sullivan, Michael E Goddard, Peter M Visscher, Jian Yang, Naomi R Wray

## Abstract

Substantial genetic liability is shared across psychiatric disorders but less is known about risk variants that are specific to a given disorder. We used multi-trait conditional and joint analysis (mtCOJO) to adjust GWAS summary statistics of one disorder for the effects of genetically correlated traits to identify putative disorder-specific SNP associations. We applied mtCOJO to summary statistics for five psychiatric disorders from the Psychiatric Genomics Consortium – schizophrenia (SCZ), bipolar disorder (BIP), major depression (MD), attention-deficit hyperactivity disorder (ADHD) and autism (AUT). Most genom-wide significant variants for these disorders had evidence of pleiotropy (i.e., impact on multiple psychiatric disorders) and hence have reduced mtCOJO conditional effect sizes. However, subsets of genome-wide significant variants had larger conditional effect sizes consistent with disorder-specific effects: 15 of 130 genome-wide significant variants for schizophrenia, 5 of 40 for major depression, 3 of 11 for ADHD and 1 of 2 for autism. In addition, we identified a number of variants that approached genome-wide significance in the original GWAS and have larger conditional effect sizes after conditioning on the other disorders. We show that decreased expression of *VPS29* in the brain may increase risk to SCZ only and increased expression of *CSE1L* is associated with SCZ and MD, but not with BIP. Likewise, decreased expression of *PCDHA7* in the brain is linked to increased risk of MD but decreased risk of SCZ and BIP.

## Introduction

Pervasive sharing of genetic risk factors between common psychiatric disorders (i.e. pleiotropy) has now been unequivocably demonstrated from genome-wide association studies (GWAS), as quantified by estimates of genetic correlation (*r_g_*)^1, 2^. The *r_g_* estimates are highest between schizophrenia and bipolar disorder (0.67, standard error (s.e.) = 0.03) but are > 0.15 for any combination of the five common disorders of schizophrenia (SCZ), bipolar disorder (BIP), ADHD, Major Depression (MD) and autism spectrum disorders (AUT) ^2, 3^. Cross-diagnosis analyses can leverage power to identify genetic risk loci shared across classical diagnostic boundaries ^4^ and can increase power for risk prediction of disorders in independent samples ^5, 6^. The shared genetic basis for psychiatric disorders contributes to an evidence base supporting a trans-diagnostic approach in clinical practice ^7^. Nonetheless, traditional diagnostic classes reflect real symptom differences at patient presentation even though it can be difficult to classify some individuals given a high-degree of concurrent and longitudinal comordibity. Since *r_g_* estimates are higher between data sets of the same disorder than between data sets of different disorders ^4, 8^ it implies some real biological basis to the classical diagnostic classes. Hence, a key question of importance in psychiatry is identification of genetic factors that are disorder specific rather than those shared across classical diagnostic groupings. Identifying such variants could aid in understanding the biological pathways that underlie the constellation of symptoms seen in each disorder.

Here, we sought to identify SNPs with disorder-specific effects by using independently collected genome-wide association study (GWAS) summary statistics for different disorders (thereby maximising contributing sample sizes). We conditioned the effect of SNPs estimated for one disorder on those of other disorders using multi-trait, conditional and joint analysis (mtCOJO)^9^, a summary-statistics based method that accounts for overlap in samples contributing to the disorder specific GWAS. We report results from conditional analyses of 5 psychiatric disorders: SCZ, BIP, MD, ADHD and AUT using association summary statistics from meta-analyses conducted by the Psychiatric Genomics Consortium (PGC) including data from 23andMe. Each disorder is conditioned on the other four disorders in one model.

## Methods

We applied the mtCOJO method as described in Zhu et al. ^9^. This method approximates a conditional analysis where the effect of a SNP on a disease is conditioned upon the covariates of the disease, but only requires summary statistics as input. As an example, if we are interested in estimating the effect of a SNP (*z*) on risk to schizophrenia (*y*) accounting for the effect of a covarying factor such as bipolar (*x*), we condition upon the effect of bipolar on schizophrenia 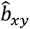, as estimated using Generalised Summary-based Mendelian Randomisation (GSMR). This can be extended to condition upon multiple covarying diseases so that the effect of the SNP on risk on the disorder of interest is estimated conditional upon the covariates on the disorder (see **Supplementary Material** for detailed description of the method).

To identify independent genome-wide significant SNPs for use as genetic instruments in mtCOJO analysis, each dataset was clumped to select independent genome-wide significant (GWS) SNPs (p < 5 × 10^-8^) using 7,762 unrelated individuals from the Atherosclerosis Risk In Community (ARIC) dataset^10^, imputed to 1000Genomes Phase III as an LD reference sample. GWS SNPs more than 1MB apart or with an r^2^ value < 0.05 were considered to be independent. GSMR accounts for any remaining LD between instruments. GSMR analysis with filtering to remove SNPs with outlier pleiotropic effects (compared to other GWS SNPs) using the HEIDI test ^11^ was performed with each disorder included both as an exposure and an outcome in combination with the other disorders. Owing to having fewer than 10 independent GWS SNPs, independent SNPs significant at p < 10^-7^ were used for GSMR analysis with autism as the exposure variable. In order to compare the estimated effects of one disorder on another from MR, we derived a conversion of the estimated effects from GSMR to the liability scale (see **Supplementary Material**).

We performed mtCOJO analysis (implemented in GCTA^12^ (http://cnsgenomics.com/software/gcta/#mtCOJO) of 5 genetically correlated psychiatric disorders using the results from large genome-wide association studies from the Psychiatric GWAS consortium (**Table 1**)^8, 13–16^, running the analysis in turn with each disorder as the outcome with the other disorders as covariates. A total of 5,275,400 SNPs with matching alleles that were in common across the 5 disorders were used for further analysis. Indels were excluded from the analysis.

**Table 1.** Summary of datasets used and results from conditional GWAS analysis

| Disorder | Cases | Controls | No. SNPs in original study | No GWS SNPs in merged data | Study reference | No. GWS SNPs in Published Study | Assumed Lifetime Disease Risk | GWS loci in conditional GWAS | New GWS loci | GWS SNPs from unadjusted GWAS with larger conditional effect size |
| --- | --- | --- | --- | --- | --- | --- | --- | --- | --- | --- |
| **SCZ** | 40,675 | 64,643 | 5,471,613 | 130 | Pardinas et al 2018 | 145 | 0.01 | 43 | 10 | 15 |
| **BIP** | 20,352 | 31,358 | 9,498,970 | 16 | Stahl et al 2018 BioRxiv | 19 | 0.01 | 4 | 3 | 0 |
| **MD** | 135,458 | 344,901 | 10,468,943 | 40 | Wray et al 2018 | 44 | 0.15 | 15 | 1 | 5 |
| **ADHD** | 19,099 | 34,194 | 6,755,648 | 11 | Demontis et al 2019 | 12 | 0.05 | 5 | 2 | 3 |
| **AUT** | 18,381 | 27,969 | 7,539,669 | 2 | Grove et al 2019 | 3 | 0.01 | 1 | 0 | 1 |

For each disorder, SNP effects conditional upon the other disorders were calculated. Results were uploaded to FUMA for annotation ^17^. Ranking SNPs according to the difference between the marginal and conditional effect sizes for each disorder is not necessarily meaningful because, for example, a SNP that has a low estimated marginal effect, so no effect on the outcome trait, will have a large conditional effect if the SNP has a large effect on the covariate traits. For the purposes of identifying which SNPs show evidence of disorder-specificity, we focus on presenting results for SNPs that were GWS with the outcome disorder in the original GWAS.

### MAGMA gene-set analysis

MAGMA gene-set analysis ^18^ as implemented in FUMA was used to investigate which sets of biologically related genes show the strongest evidence of association in the conditional analyses.

### Genetic Correlation

LD-score regression ^19^ was used to estimate the genetic correlation between the conditional and unadjusted GWAS results.

### Summary Mendelian Randomisation

To investigate the potential functional relevance of SNPs with disorder-specific effects, we applied the SMR approach ^11^, integrating eQTL (SNP-gene expression association) and mQTL (SNP_DNA methylation association) to the results from the conditional analyses. eQTL data from brain tissue were derived from a meta-analysis of the GTEx study, the Common Mind Consortium (CMC) and the Religious Orders Study and Memory and Aging Project (ROSMAP). The details of the meta-analysis have been described elsewhere^20^. Using meta-analysis results across brain tissues and studies is justified owing to the high correlation in effect sizes between tissues^20^. Only genes with a cis-eQTL with p_eQTL_ < 5 × 10^-8^ were included in the analysis. Experiment-wide significance accounting for testing multiple SNPs across multiple traits was set at p_SMR_ = 1.9 × 10^-06^ and the threshold for no evidence of heterogeneity due to pleiotropy at p_HEIDI_ > 0.01. Individual-level genotypes from the ARIC data (n = 7,762 unrelated individuals) ^10^ were used to estimate LD for the HEIDI test.

To test for the effects of disorder-specific variants on DNA methylation, we used SMR to integrating trait association data with meta-analysed brain mQTL data set from Jaffe et al. (n = 526) ROSMAP (n = 486) and fetal brain mQTL data from Hannon et al ^21^. Only probes with at least one cis-mQTL with p < 5 × 10^-8^ were included in the SMR analysis. Probes that passed the significance threshold of 1.56 × 10^-7^ and did not show evidence of heterogeneity as indicated by the HEIDI test were considered to be significant.

### Cell-type specificity for disorders

To gain insight into the cell types that are important for each disorder, we evaluated whether genes associated with specific brain cell-types are enriched for association with each of the disorders. Using data from single-cell sequencing experiments in mice, the cell-type specificity of each gene was calculated by comparing the expression of a gene in a given cell-type to that across all cell types ^22^. MAGMA was used to calculate gene-based association statistics and to evaluate whether genes with high specificity in a given cell-type are enriched for association with a disorder. The enrichment analysis was performed for both unadjusted and conditional GWAS for all 5 disorders. To investigate whether there was a significant change in the cell-type enrichment after conditioning, MAGMA analysis was performed using the enrichment Z-scores from the unadjusted GWAS as covariates in the analysis and a conditional enrichment for all level 1 cell types analysed in Skene et al. ^22^ was estimated.

## Results

### Baseline statistics

After merging GWAS summary statistics for the five psychiatric disorders 5,275,400 autosomal SNPs remained (**Table 1**). The number of independent genome-wide significant SNPs annotated by FUMA ^17^ is much greater for SCZ (M =130) compared to the other disorders (M =16, 40, 11, 2 for BIP, MD, ADHD, AUT respectively) reflecting mostly sample size, but also genetic architecture, and population risk. Linkage disequilibrium score regression (LDSC) estimates of SNP-based heritability on the liability scale and genetic correlations were all significantly different from zero (**Table 2**). Genetic correlations were highest between SCZ and BIP (*r_g_* = 0.67 (s.e. = 0.03)) and lowest between BIP and ADHD (*r_g_* = 0.15 (s.e. = 0.04)). The LD-score regression intercept was significantly greater than zero for the majority of pairs of disorders reflecting sample overlap in the GWAS studies. The intercept was highest between ADHD and AUT due to substantial overlap in controls.

**Table 2.** Estimated SNP-based heritability on the liability scale, genetic correlation and LD-score intercepts estimated from LD-score regression

|  | SCZ | BIP | MDD | ADHD | AUT |
| --- | --- | --- | --- | --- | --- |
| SCZ | **0.23 (0.01)** | *0.21 (0.01)* | *0.03 (0.01)* | 0.02 (0.01) | 0.008 (0.01) |
| BIP | 0.67 (0.02) | **0.19 (0.01)** | *0.05 (0.007)* | *0.03 (0.006)* | 0.009 (0.008) |
| MD | 0.36 (0.02) | 0.35 (0.02) | **0.08 (0.004)** | *0.10 (0.008)* | *0.09 (0.008)* |
| ADHD | 0.18 (0.03) | 0.15 (0.04) | 0.43 (0.03) | **0.22 (0.01)** | *0.35 (0.008)* |
| AUT | 0.23 (0.05) | 0.15 (0.05) | 0.43 (0.04) | 0.36 (0.05) | **0.12 (0.01)** |

LD-score SNP-based heritability on the liability scale and standard error reported on diagonal
$r_g$ and standard error reported below the diagonal
Bivariate ldsc intercept reported above the diagonal. Value significantly greater than zero (in italics) quantify sample overlap

The GSMR analyses highlights some asymmetries in the estimates of the causal effects of one disorder on another (**Table 3**). In particular, the estimated liability 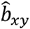 when considering MD as an exposure for each trait is higher than the estimates in the reverse direction. One explanation is that since MD is so common and is frequently comorbid with other disorders that MD samples include those diagnosed and undiagnosed with other disorders. However, if model assumptions are violated it may have greater impact when there is a large difference in lifetime risk between the pairs of disorder. However, countering this, we find a higher 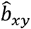 from AUT to ADHD than from ADHD to AUT, but the standard errors on estimates are much higher for these disorders. Interpretation of these 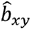 estimates depends on the nature of the shared genetic contributions to psychiatric disorders that may reflect a complex mix of types of pleiotropy, where some sets shared of variants may have more correlated effect sizes than other sets of shared variants.

**Table 3.**
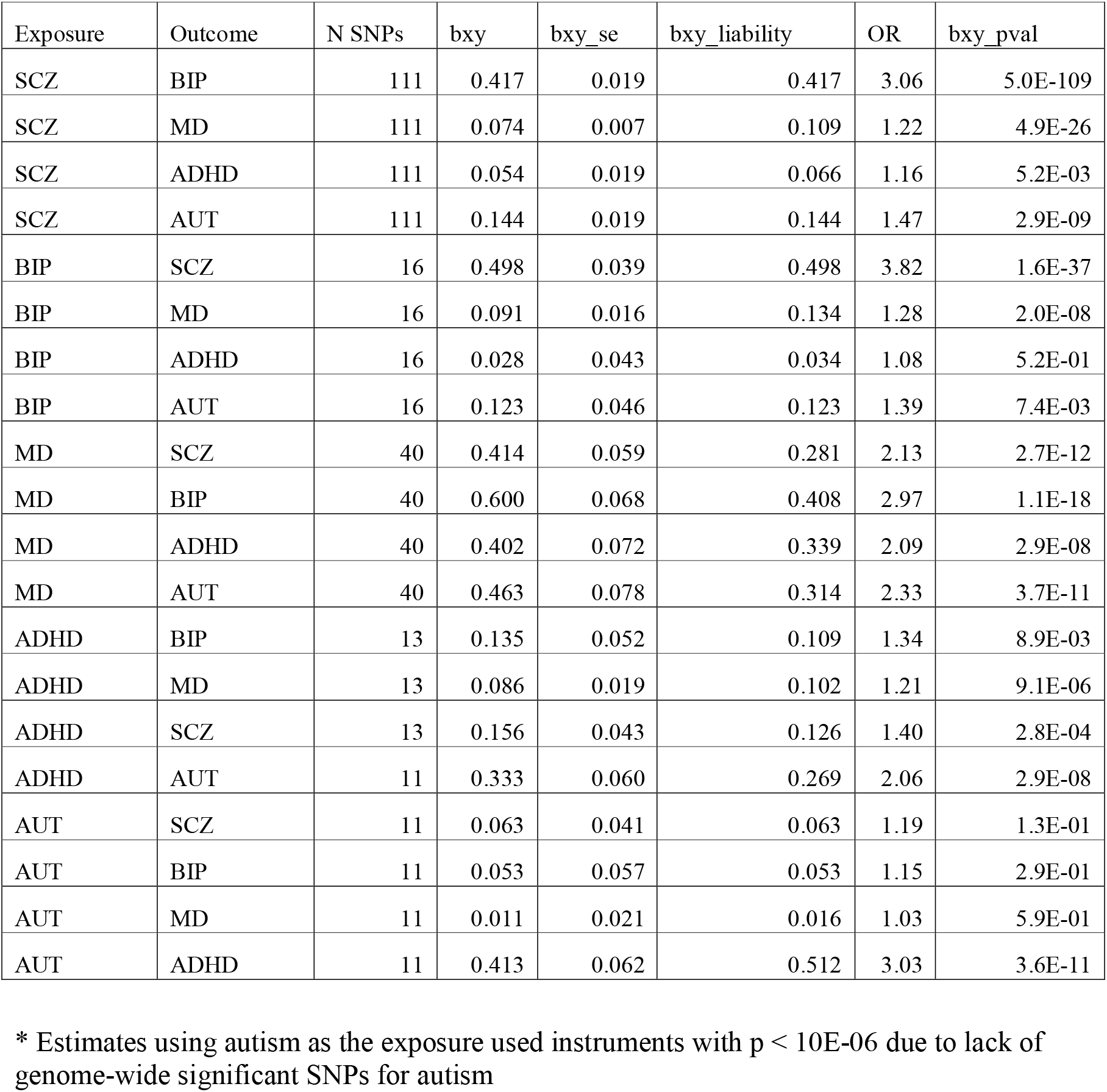
GSMR estimates of causal effect of each psychiatric disorder on the others with conversion to the log odds ratio and liability scales

### Changes in Genetic Correlation

The impact of the conditioning is demonstrated by the changes in the estimates of 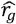 comparing original and conditional GWAS results. The 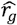 between SCZ conditional on the other disorders (denoted SCZ_cond_) and SCZ remained high at 0.93, while between SCZ_cond_ and BIP it was much reduced (from 0.67 prior to conditioning to 0.36, after conditioning). It is noted that b_zy_ is eliminated in the conditional analysis only if the SNP effect is mediated by trait *x*. Therefore, there is remaining genetic correlation because of pleiotropic SNP effects. A similar pattern of changes in genetic correlation with other traits was seen for the analyses with the other disorders as the outcome variable (**Supplementary Table 1**).

### mtCOJO genome-wide significant SNP results

As expected because of pleiotropy between disorders, conditional analysis leads to a reduction in the mean test statistic across all SNPs in the genome and hence the number of independent SNPs reaching the significance threshold (5×10^-8^) is reduced (**Table 1**). For each disorder, we present results for all independent SNPs significant in the unadjusted analysis or the conditional analysis (**Supplementary Table 2**). GWS SNPs that are more significantly associated in the conditional analysis than the unadjusted analysis are shown in **Table 4**. A larger conditional effect size suggests that these variants are disorder-specific or have heterogeneous effects across disorders.

**Table 4.** Results for SNPs that were genome-wide significant in the conditional analysis and have larger estimated conditional effect sizes than in the original GWAS

| Disorder | SNP | CH<br>R | Position | A1 | Adjusted<br>beta | Unadjusted<br>beta | SE<br>Adjusted<br>beta | Adjusted p-value | Unadjusted p-value | nearestGene |
| --- | --- | --- | --- | --- | --- | --- | --- | --- | --- | --- |
| SCZ | rs3764002 | 12 | 108618630 | C | 0.083 | 0.054 | 0.012 | 1.9E-12 | 6.1E-07 | WSCD2 |
| SCZ | rs6095357 | 20 | 47523865 | A | -0.069 | -0.048 | 0.011 | 1.2E-10 | 1.2E-06 | ARFGEF2 |
| SCZ | rs7790864 | 7 | 28478625 | A | -0.062 | -0.044 | 0.011 | 6.3E-09 | 7.2E-06 | CREB5 |
| SCZ | rs1054972 | 19 | 1852582 | A | 0.076 | 0.053 | 0.013 | 6.4E-09 | 1.3E-05 | KLF16 |
| SCZ | rs2867673 | 7 | 71752652 | T | 0.060 | 0.049 | 0.010 | 9.4E-09 | 4.1E-07 | CALN1 |
| SCZ | rs6564668 | 16 | 79457393 | C | -0.060 | -0.038 | 0.010 | 1.0E-08 | 7.9E-05 | RP11-467I7.1 |
| SCZ | rs11922765 | 3 | 95047279 | G | -0.060 | -0.044 | 0.010 | 1.2E-08 | 4.4E-06 | RPS18P6 |
| SCZ | rs2973038 | 5 | 37833781 | C | 0.066 | 0.051 | 0.012 | 1.3E-08 | 1.7E-06 | GDNF |
| SCZ | rs10903945 | 10 | 363275 | C | 0.057 | 0.040 | 0.010 | 3.1E-08 | 3.3E-05 | DIP2C |
| SCZ | rs10282935 | 8 | 38703797 | A | 0.058 | 0.041 | 0.011 | 3.9E-08 | 3.2E-05 | TACC1 |
| SCZ | rs6701877 | 1 | 174015259 | G | -0.096 | -0.073 | 0.014 | 1.5E-11 | 2.4E-08 | RP11-160H22.3 |
| SCZ | rs7372313 | 3 | 135872958 | G | -0.069 | -0.062 | 0.010 | 4.3E-11 | 1.5E-10 | MSL2 |
| SCZ | rs1765142 | 11 | 30378559 | C | 0.065 | 0.058 | 0.011 | 1.5E-09 | 1.1E-08 | ARL14EP |
| SCZ | rs55646993 | 7 | 105017864 | G | -0.062 | -0.053 | 0.010 | 2.2E-09 | 3.8E-08 | SRPK2 |
| SCZ | rs150437760 | 14 | 59981768 | A | 0.131 | 0.121 | 0.024 | 3.7E-08 | 4.6E-08 | CCDC175 |
| BIP | rs12554512 | 9 | 23352293 | T | -0.083 | -0.066 | 0.014 | 1.6E-09 | 1.3E-06 | ELAVL2 |
| BIP | rs6891181 | 5 | 80849101 | T | -0.081 | -0.075 | 0.014 | 1.5E-08 | 1.3E-07 | SSBP2 |
| BIP | rs12268910 | 10 | 111878510 | T | -0.097 | -0.091 | 0.018 | 3.3E-08 | 2.7E-07 | ADD3 |
| MD | rs11697370 | 20 | 47731767 | T | -0.031 | -0.023 | 0.005 | 3.3E-09 | 3.5E-06 | STAU1 |
| MD | rs27732 | 5 | 87992576 | A | 0.034 | 0.031 | 0.005 | 1.2E-11 | 1.9E-10 | MEF2C |
| MD | rs1806153 | 11 | 31850105 | T | 0.037 | 0.036 | 0.006 | 8.8E-10 | 1.2E-09 | RCN1 |
| MD | rs1354115 | 9 | 2983774 | A | 0.029 | 0.028 | 0.005 | 1.7E-08 | 2.4E-08 | CARM1P1 |
| MD | rs301799 | 1 | 8489302 | T | -0.028 | -0.026 | 0.005 | 2.5E-08 | 4.7E-08 | RERE |
| ADHD | rs78648104 | 6 | 50683009 | T | 0.136 | 0.124 | 0.023 | 4.3E-09 | 3.6E-07 | TFAP2D |
| ADHD | rs2244336 | 10 | 8831827 | C | 0.071 | 0.069 | 0.013 | 3.8E-08 | 3.7E-07 | ENSG00000270234 |
| ADHD | rs12410444 | 1 | 44188719 | A | 0.107 | 0.106 | 0.014 | 4.2E-15 | 3.8E-13 | ST3GAL3 |
| ADHD | rs13023832 | 2 | 215219808 | A | 0.133 | 0.117 | 0.020 | 1.2E-11 | 1.6E-08 | SPAG16 |
| ADHD | rs281320 | 15 | 47769424 | T | -0.080 | -0.074 | 0.013 | 1.8E-10 | 3.1E-08 | SEMA6D |
| AUT | rs10099100 | 8 | 10576775 | C | 0.084 | 0.084 | 0.014 | 1.2E-09 | 1.0E-08 | SOX7 |

Given that SCZ is the disorder with the largest number of significant SNPs, we focus mostly on the results from the SCZ conditional analysis. Of the 130 SNPs from the unadjusted SCZ GWAS, five were more significant after adjusting for the other disorders (all of which had opposite direction of effects for BIP – **Supplementary Table 2**) and a further eight had a larger estimated effect size after conditioning. Forest plots for the four most significant SCZ SNPs from the conditional analysis (two of which were associated p<5×10^-8^ in the unadjusted analysis) are shown in **Figure 1**.

**Figure 1.**
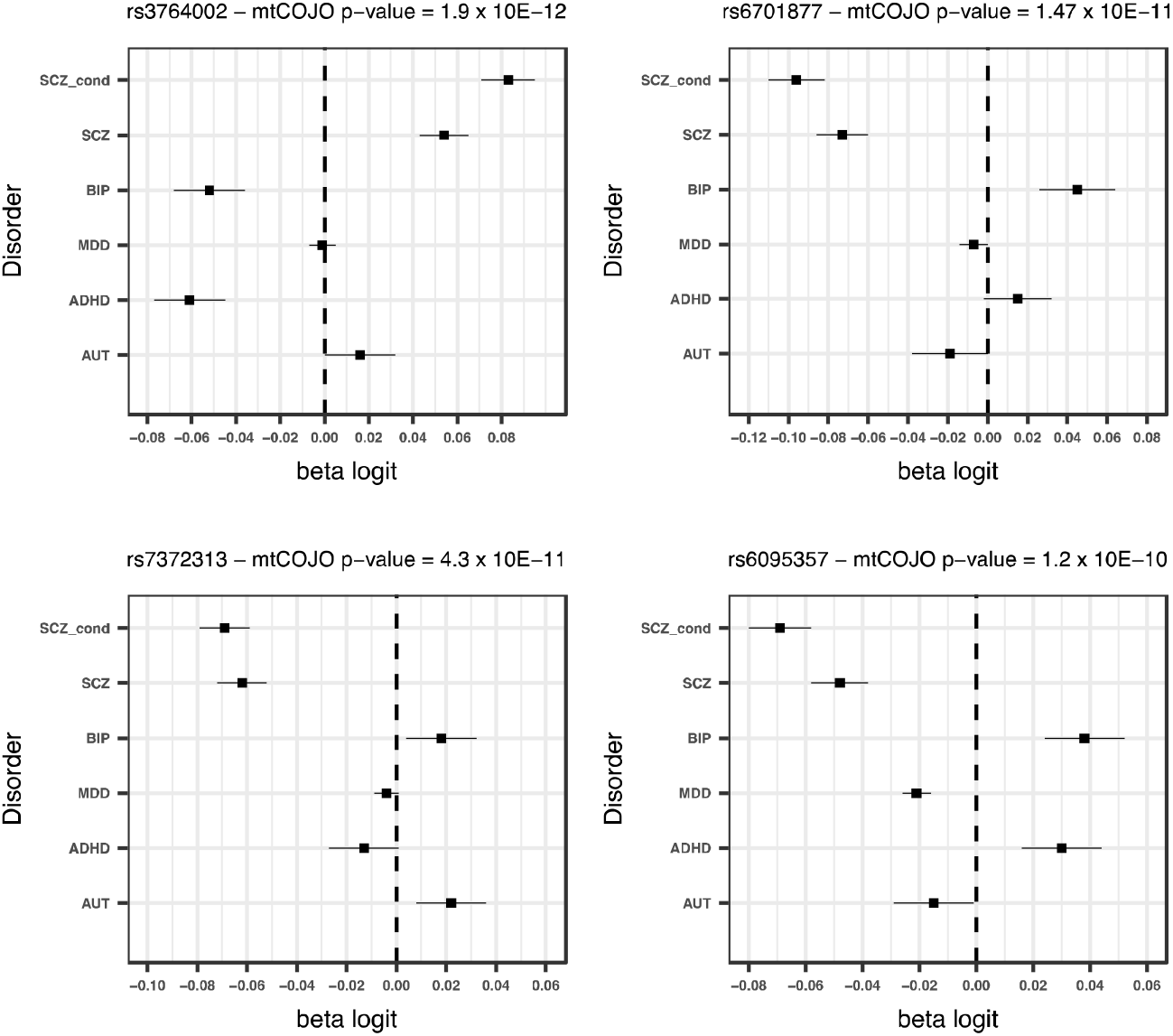
Forest plots for the 4 most significant SNPs in SCZ mtCOJO analysis with larger conditional effect sizes

For all disorders except for AUT, a number of SNPs surpass the significance threshold that were not significant in the original GWAS. For schizophrenia, ten SNPs that were significant in the conditional analysis and not in the original GWAS (**Table 4**). All 10 SNPs have opposite effects for BIP, so that the allele that predisposes to SCZ is in the protective direction for BIP. Although these opposite effects could be due to ascertainment, among them are variants in or near genes with annotated biological functions that are potentially relevant for SCZ. For instance a SNP that was significant in the conditional analysis (rs2973038 – p_adj_ = 1.28 × 10^-08^; p_scz_ = 1.72 × 10^-06^) is located in the Glial Cell Derived Neurotrophic Factor (*GDNF*), a gene that encodes a protein that enhances the survival of midbrain dopaminergic neurons ^23^, and is expressed during development ^24^.

All SNPs that were associated with BIP at p< 5×10^-8^ in the original GWAS were less significant in the conditional analysis, showing evidence that they have some pleiotropic effect across disorders. Notably, this includes genes involved in calcium signalling, dopaminergic signalling and synaptic plasticity, indicating these processes may be important across psychiatric disorders. Three SNPs that were not significant in the BIP GWAS were significant in the conditional analysis (**Table 4, Supplementary Table 2, Supplementary Figure 2**).

For each of the remaining disorders (MD, ADHD and AUT), we found that a small proportion of the existing significant SNPs had larger conditional effect sizes and one MD SNP and two ADHD SNPs that were not significant in the original GWAS became significant after conditioning (**Table 4, Supplementary Table 2**). Forest plots for significant SNPs that had increased conditional effect sizes are shown in **Supplementary Figures 3-5**

### SMR analysis

Changes in the expression of 9 genes were significantly associated with the 5 disorders (0 for BIP, 5 for SCZ, 3 for MD and 1 for ADHD, 0 for AUT) after conditioning and removal of genes in the MHC (**Supplementary Tables 3-4**), and a total of 72 DNA methylation sites (2 for BIP, 18 for SCZ, 37 for MD, 8 for ADHD, and 6 for AUT) were significantly associated with the 5 conditional traits (**Supplementary Table 3-4**).

Significant SMR results for gene expression where the associated SNP is more significant in the conditional analysis are presented in **Supplementary Table 3**. Three out of 5 significant SMR associations for SCZ were with SNPs where the conditional significance was greater than in the unadjusted analysis. One SNP - rs3759384 – is associated with decreased expression of *VPS29* in the brain and significantly increased risk for SCZ in the unadjusted analysis and has a larger conditional effect size (Supplementary Figure 6), indicating that *VPS29* may be linked to the development of SCZ and not other disorders. The VPS29 protein is a component of the retromer complex which prevents the degradation of certain proteins including signalling receptors, ion channels and small molecule transporters. The complex is essential for maintenance of neurons and has been implicated in the etiology of a number of neurodegenerative disorders ^25^.

One of the three associations for MD was with a SNP (rs7732179) with greater significance in the conditional analysis. The same variant shows evidence of association with SCZ but with opposite directions of effect (b_SCZ_ = −0.045; p_SCZ_ = 1.7 × 10^-6^ and b_BIP_ = −0.029; p_BIP_ = 0.027). The A allele confers risk to MDD but is protective for SCZ and BIP (Supplementary Figure 7). The SNP is associated with expression of *PCDHA7* in the brain. This gene encodes a member of the protocadherin family of genes located together on chromosome 5. A significant association was also found in this region in the DNA methylation analysis of MD. Little is known about the exact function of these genes, however they are concentrated at the synaptic junction suggesting a key role in neuronal signalling^26^.

Out of 72 significant DNA methylation sites, 34 were associated with SNPs with higher significance in the conditional analyses (1 for BIP, 3 for SCZ, 21 for MD, 4 for ADHD and 5 for AUT) (**Supplementary Table 3**). It is noteworthy that one variant (rs2064853) was significantly associated with both SCZ and MD and DNA methylation near the *CSE1L* gene, but with opposite alleles increasing risk to each disorder (Supplementary Figure 8).

We investigated whether genes identified in the gene expression SMR or that are the closest gene to a significant methylation site are the primary target for FDA-approved drugs. We identified two genes that are targeted by medications. The serotonin receptor gene *HTR1D* which was identified in the DNA methylation analysis for MD is the primary target of the migraine drug naratriptan. Individuals with migraine are at 2-4 fold higher risk of developing depression and these results may suggest that triptans, used to treat migraines, could also be effective for MD.

The second drug target identified is with *MPL* and ADHD. This gene is targeted by romiplostim, an orphan drug developed for treatment of chronic idiopathic thrombocytopenic purpura.

### MAGMA gene-set analyses

We conducted MAGMA gene-set analysis in FUMA to identify pathways and gene-sets that are enriched for association with the disorders after conditional analyses and to identify which sets become more or less significant after conditioning. Results for each disorder are presented in **Supplementary Table 5**. After conservative Bonferroni correction for the number of gene-sets tested for each disorder, three gene sets were significant - two for SCZ conditional analysis and one for AUT. For SCZ, the two significant sets were go:establishment of localization in the cell and GO:Dendrite, of which establishment of localization had a more significant p-value in the conditional analysis (**Supplementary Table 5**). For AUT, the gene-set GO:Dendrite_morphogenesis was significant after multiple testing and had a more significant p-value in the conditional analysis, potentially implicating genes expressed in dendrites in autism-specific pathology.

### Cell-type specificity for disorders

The results from the cell-type enrichment analyses of raw and conditional analyses are shown in **Figure 2**. Consistent with previous results, the original SCZ results were enriched in medium spiny neurons (MSNs), pyramidal CA1 cells, pyramidal SS1 cells, interneurons and serotonergic neurons (**Supplementary Table 6**). All of these cell types also show some evidence of association with BIP and to a lesser extent MD, consistent with the genetic correlation between disorders and hence show reduced enrichment in the SCZ conditional analysis. All enriched cell-types for SCZ remained significant after conditioning except for serotonergic neurons, indicating that genes specific to this cell-type may increase risk to all five disorders. Enrichment in interneurons was found for SCZ, BIP and MDD indicating their potential importance across all 3 disorders. After conditioning, this cell-type was still significantly enriched in SCZ and MDD, but not BIP. This may reflect that the sample size of the BIP analysis is smaller than for SCZ and MDD.

**Figure 2.**
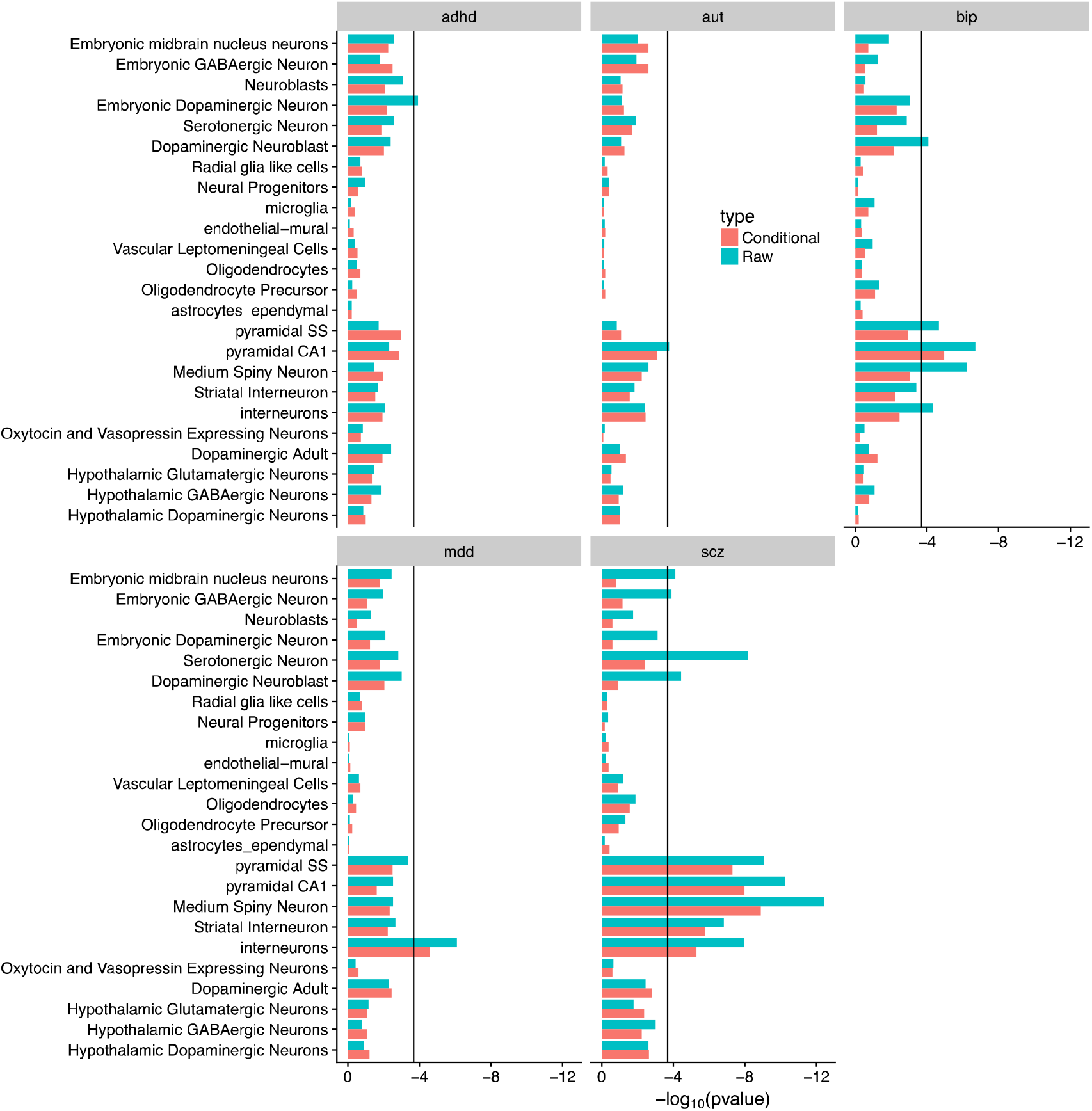
Results from brain cell-type enrichment analyses of raw and conditional GWAS analyses

## Discussion

Our goal was to identify genetic variants that show disorder specific association by conducting a summary statistics based GWAS analysis for each of five psychiatric disorders conditioning on GWAS results from the other disorders. As expected, given the high degree of pleiotropy across disorders, compared to original GWAS results the number of SNPs associated at the threshold of genome-wide significance is very much reduced for each conditional GWAS. The conditional analyses identified SNPs that increased in significance compared to the unconditional results, which reflects SNPs which have opposite effect size in one or more of the disorders that are included in the conditional analysis. A small number of SNPs that were not previously significant surpassed the p<5×10^-8^ threshold in the conditional analyses. Since effect sizes are estimated with error, if these SNP associations are true positives we would expect them to become genome-wide signifcant in future disorder specific GWAS that are larger and more powered. For SCZ and MD, the proportion of previously genome-wide significant SNPs that have larger conditional effect sizes is approximately 10%. The conditional results for BIP, ADHD and AUT were less insightful, as expected, since the GWAS of these disorders have identified fewer significant loci. Future larger samples are needed to exclude results that have achieved increased statistical significance in conditional analyses through chance. We utilise mtCOJO as a method that uses summary statistics to quickly screen for SNP associations. Functional annotation can help prioritise the associations of most interest. It will be important to understand why a variant increases risk only to that disorder and not to others.

By integrating conditional GWAS results with SNP-gene expression and SNP-methylation results, we identify decreased expression of *VPS29* as a potential biological mechanism underlying schizophrenia. The variant that increases risk to SCZ and is associated with decreased expression of VPS29 in brain tissue shows no evidence for association with other psychiatric disorders. Moreover, SNPs associated with decreased expression of *PCDHA7* and decreased methylation near other members of the protocadherin gene family on chromosome 5 may increase risk of MD, but be protective for SCZ and BIP. Methylation in the promoter of the CSE1L gene, whose encoded protein influences cellular proliferation and has been linked to progression of a number of cancers, shows evidence of increasing risk to SCZ but being protective for MD.

Consistent with the large degree of pleiotropy between disorders, we found that most of the significant biological pathways for each disorder had reduced significance after conditioning. Pathway analysis of conditional results identified a potential role for genes expressed in dendrites in both autism and schizophrenia. Likewise, for the cell-type enrichment analysis, there was a reduction in the enrichment for most cell types in each disorder after conditioning. For SCZ, the previously identified enrichments in pyramidal SS1 and CA1 cells as well as medium spiny neurons remained significant after conditioning, despite also showing evidence for enrichment in BIP. The largest change in enrichment was for serotonergic neurons, indicating that genes highly expressed there are important across all psychiatric disorders.

We provide an analysis framework for conditional cross-disorder analyses using summary statistics. Our study was motivated to improve on the SCZ case vs. BIP case analyses that utilised PGC cohorts for which both SCZ and BIP genotyped samples were available ^27^, but which necessarily excluded 28% of cases that could not be allocated into matched cohorts. They identified 5 SNPs associated at p<5×10^-8^. We conducted an analysis of SCZ conditional on BIP and performed a lookup of those SNPs in the unadjusted and adjusted results. All but one (rs200005157) of their associated SNPs were matched directly or to an LD proxy (**Supplementary Table 7**). All show increased statistical significance in the conditional analysis. We identified more disorder-specific SNPs (10 specific to SCZ) consistent with the larger sample sizes afforded from using summary statistics, highlighting that mtCOJO is an efficient method for screening for disorder-specific SNPs for two or more related disorders. An in depth discussion of the mtCOJO method and potential limitations are given in the Supplementary Material.

In conclusion, our results suggest that mtCOJO is an efficient method for identifying variants with disorder-specific effects and they represent a small fraction of variants identified for each disorder to date, reflecting the high degree of pleiotropy between disorders. Nonetheless, we identify several loci that have evidence of being disorder-specific. Further research in human studies should focus on whether the disorder-specific variants associate with specific symptoms in mixed clinical populations.

## Acknowledgements

This work is supported by grants from the National Health and Medical Research Council of Australia (1087889, 1145645, 1113400, 1078901,1078037), and the Sylvia & Charles Viertel Charitable Foundation. The PGC has received major funding from the US National Institute of Mental Health and the US National Institute of Drug Abuse (U01 MH109528 and U01 MH1095320). We thank the research participants and employees of 23andMe, Inc. for contributing to this study. This paper would not have been possible without the generosity of participants in the many studies that comprise the final meta-analyses and the dedication of many clinicians and research staff who have collected the data and made them publically available. Acknowledgments for specific datasets are provided in the Supplementary Material.

