## Supplementary Methods and Figures for "Conditional GWAS analysis identifies putative disorder-specific SNPs for psychiatric disorders"

**Table of Contents**

1. **Discussion of mtCOJO method for conditional analysis (**Page 1)
2. **Summary of estimates of *b_xy_* for all combinations of quantitative and binary traits** (Page 1)
3. **Simulations to confirm derivations** (Page 3)
4. **Supplementary Figures (**Page 7**)**
5. **Acknowledgments** (Page 15)

**1. Discussion of mtCOJO method for conditional GWAS analysis**

**Multi-trait, conditional and joint analysis (mtCOJO)**

We used the mtCOJO methodology^9^ to conduct a conditional GWAS analysis of psychiatric disorders. mtCOJO was developed to generate GWAS summary statistics results for a target trait conditional on a covariate trait, recognising that the covariate trait is frequently not recorded in the individuals measured for the target trait. The mtCOJO method allows conditioning on the exposure by borrowing information from independently collected data, with the data linked through their SNP associations. Briefly, for two traits (*y* and *x*) with association effects for SNP z ( $\hat{b}_{zy}$ and $\hat{b}_{zx}$respectively) the association effect estimates for trait *y* conditional on trait *x* is estimated as $\hat{b}_{zy}|\hat{b}_{xy}=\hat{b}_{zy}-\hat{b}_{zx}\hat{b}_{xy}$ (see Zhu et al. ^9^ for details), where $\hat{b}_{xy}$ is the effect of trait *x* on trait *y*, as estimated in generalised summary-based Mendelian randomisation (GSMR) analyses. If exposure trait *x* is causal of the outcome trait *y* or the two traits are pleiotropically related, we expect the genetic correlation estimated from the conditional GWAS result for trait *y* and that for trait *x* to be reduced by a factor proportional to the estimated effect of *x* on *y*. In a special case that if traits *x* and *y* are genetically identical we expect this correlation to be zero.

Classical MR analyses have the hypothesis of causality of an exposure trait on an outcome trait. Here, when the two disease traits are psychiatric disorders, we have no hypothesis of causality, but rather use GSMR to demonstrate significant bi-directional association of SNPs that are GWS in one disorder with those in another disorder and to estimate the weights for mtCOJO.

mtCOJO allows the analysis conditioning on multiple covarying diseases, so that the effect of a SNP on risk on the disorder of interest conditional upon the covariates on the disorder, is given by $\hat{b}_{zy}|{\hat{\mathbf{b}}}_{xy}=\hat{b}_{zy}-{\hat{\mathbf{b}}}_{zx}^{t}{\hat{\mathbf{b}}}_{xy}$ where $\hat{b}_{zy}$ is the SNP effect on the disease, ${\hat{\mathbf{b}}}_{xy}$ is a *t*-length vector with the *i*-th element $\hat{b}_{x_{i}y}$ being the effect of $x_{i}$ on the disease when all the covariates are fitted jointly, and${\hat{\mathbf{b}}}_{zx}$ is a *t*-length vector of SNP effects on **x.** The method is robust to sample overlap between studies.

If the genetic correlation between two disorders is not driven by causality but a large number of pleiotropic effects (generally in the same direction) because of shared genetic pathways, then the GSMR $\hat{b}_{xy}$reflects an average pleiotropic association between the two disorders. Nonetheless, conditioning on the average effect has meaning, and our conditional results can be used in pathway analyses to identify functionally relevant pathways that are associated with a disorder after accounting for effects of covarying disorders.

While it is possible that differences in quality control analysis or calling/imputation of SNPs across studies could lead to heterogeneity in the results and thus significant effects in the conditional analysis, these datasets were generated using the same analysis pipeline developed by the Psychiatric GWAS Consortium, and hence the likelihood of such heterogeneity is reduced.

**2. Summary of estimates of *b_xy_* for all combinations of quantitative and binary traits**

When estimating $b_{xy}$ using GSMR in instances where the exposure trait and outcome trait are quantitative traits and the effect sizes have been standardised, the effect estimate is directly interpretable as SD units of y per SD unit of x. When using disease traits as exposures, $b_{xy}$ is estimated on the logit scale and used in mtCOJO. The estimates on the logit scale can be difficult to interpret when disorders have a different prevalence and the proportion of cases and controls used in the GWAS is different from the population prevalence We derive transformations to make the$b_{xy}$estimates interpretable when the exposure trait is a disease.

The key transformations of the $b_{xy}$ on the logit scale to the logOR and liability scale are

$$\hat{b}_{xy(\mathrm{logOR})}=\frac{z}{K(1-K)}\hat{b}_{xy(logit)}$$

$$\hat{b}_{xy(liab)}=\frac{z_{K(x)}K_{y}(1-K_{y})}{z_{K(y)}K_{x}(1-K_{x})}\hat{b}_{xy(logit)}$$

where *K_x_* is the lifetime risk of the exposure trait and *K_y_* is the lifetime risk of the outcome trait and $z_{K}$ is the height of the normal distribution at the truncation point on the liability scale corresponding to risk *K*. $\hat{b}_{xy(logOR)}$can be interpreted as the log of the odds ratio of *y* per 1 SD increase in liability in *x*. The estimate on the liability scale can be interpreted as the increase in liability in to disorder *y* per 1 SD increase in liability to *x*. Note that when two disorders have the same assumed prevalence, as is the case with schizophrenia and bipolar, $\hat{b}_{xy(logit)}= \hat{b}_{xy(liab)}$ . See the derivation below.

For two traits (*y* and *x*) with association effect sizes for SNP z ($\hat{b}_{zx}$ and $\hat{b}_{zy}$, respectively) the association effect size estimates for trait *y* conditional on trait *x* ($\hat{b}_{zy}|x$) is estimated as $\hat{b}_{zy}|x\approx{\hat{b}_{zy}|{\hat{\mathbf{b}}}_{xy}= \hat{b}}_{zy}-\hat{b}_{zx}\hat{b}_{xy}$ , where ${\hat{\mathbf{b}}}_{xy}=\{ \hat{b}_{xy\left( 1 \right)}, \hat{b}_{xy\left( 2 \right)}, \cdots,\hat{b}_{xy(m)}\}$ with $\hat{b}_{xy(i)}=\hat{b}_{zy(i)}/\hat{b}_{zx(i)}$, and ${\hat{\mathbf{b}}}_{xy}\sim N(\boldsymbol{1}b_{xy},\mathbf{V})$ where **1** is an *m*×1 vector of ones and **V** is the variance-covariance matrix of ${\hat{\mathbf{b}}}_{xy}$, and *m* is the number of genome-wide significant SNPs selected as independently associated with trait *x.* $\hat{b}_{xy}$ is the effect of trait *x* on trait *y*, which can be calculated through GSMR analysis, $\hat{b}_{xy}$ = ${(\mathbf{1}^{'}\mathbf{V}^{-1}\mathbf{1})}^{-1}\mathbf{1}^{'}\mathbf{V}^{-1}{\hat{\mathbf{b}}}_{xy}$ (1).

When trait *x* is a dichotomous trait we denote the population lifetime risk as *K_x_*, $v_{K_{x}}$ as height of the normal curve when truncated at *K_x_*, and the proportion of cases in the samples used to generate the SNP effects as *P_x_*. The effect size estimated from logistic regression of *x* on SNP *z* is on the logit scale $\hat{b}_{zx(logit(x))}$, such that exp($\hat{b}_{zx(logit(x))}$) is the odds ratio of the SNP for cases of disease *x*. From Lloyd-Jones et al(2), the relationship between $\hat{b}_{zx(logit(x))}$and $\hat{b}_{zx}$ on the observed scale from a case-control sample with proportion of cases $P_{x}$ ($\hat{b}_{zx(01[P_{x}])}$) is

$b_{zx(01[P_{x}])}\approx P_{x}(1-P_{x})b_{zx(logit(x))}$

From Lee et al. 2011 (3) and Dempster and Lerner(4) the relationship between $\hat{b}_{zx}$ on the liability scale and ( $\hat{b}_{zx(liab)}$) and $\hat{b}_{zx(01[P_{x}])}$ is

$\hat{b}_{zx(liab)}\boldsymbol{=}\frac{K_{x}\left( 1-K_{x} \right)}{v_{K_{x}}P_{x}(1-P_{x})}\hat{b}_{zx\left( 01[P_{x}] \right)}$.

Hence $\hat{b}_{zx(liab)}\boldsymbol{=}\frac{K_{x}\left( 1-K_{x} \right)}{v_{K_{x}}}\hat{b}_{zx\left( logit(x) \right)}$.

So when *y* is a quantitative trait and *x* binary, the $\hat{b}_{xy}$ calculated from available summary statistics is $\frac{\hat{b}_{zy}}{\hat{b}_{zx\left( logit(x) \right)}}=\hat{b}_{xy(y:logit(x))}$. A more interpretable regression coefficient is $\hat{b}_{xy(liab)}\boldsymbol{=} \frac{\hat{b}_{zy}}{\hat{b}_{zx\left( liab \right)}}= \frac{v_{K_{x}}K_{y}}{K_{x}(1-K_{x})}\hat{b}_{xy(y:logit(x))}$. Similarly, if *y* is a dichotomous phenotype, the $\hat{b}_{xy}$ calculated from available summary statistics is $\frac{\hat{b}_{zy(logit\left( y \right))}}{\hat{b}_{zx\left( logit(x) \right)}}=\hat{b}_{xy(logit(y):logit(x))}$, a more interpretable regression coefficient is either as logit of *y* per phenotypic liability SD unit of *x*:$\hat{b}_{xy(logit\left( y \right):liab(x))}=\frac{v_{K(x)}}{K_{x}(1-K_{x})}\hat{b}_{xy(logit\left( y \right):logit\left( x \right))}$ , or as phenotypic liability SD unit of *y* per phenotypic liability SD unit of *x*, $\hat{b}_{xy(liab\left( y \right):liab(x))}=\frac{\hat{b}_{zy\left( liab \right)}}{\hat{b}_{zx\left( liab \right)}}=\frac{v_{K(x)}K_{y}(1-K_{y})}{v_{K(y)}K_{x}(1-K_{x})}\hat{b}_{xy(logit\left( y \right):logit\left( x \right))}$. So, for schizophrenia and bipolar disorder since we assume $K_{x}= K_{y}$ = 0.01, then $\hat{b}_{xy(liab\left( y \right):liab(x))}=\hat{b}_{xy(logit\left( y \right):logit\left( x \right))}$. The interpretation and scaling of $\hat{b}_{xy}$ for any combination of binary and quantitative traits is summarised in **Supplementary Table 9**.

**Supplementary Table 9.**

| a) Units of $\hat{b}_{xy}$  given $\hat{b}_{xy(i)}=\hat{b}_{zy(i)}/\hat{b}_{zx(i)}$  b) Scale of estimation | **Trait y is quantitative**  **Linear regression GWAS**  $\hat{b}_{zy}$**:** Allelic effect in phenotypic SD units of *y* | **Trait y is binary disease**  **Logistic regression GWAS**  $\hat{b}_{zy(logit(y))}$**:** Allelic effect in ln(OR) of *y* |
| --- | --- | --- |
| **Trait x is quantitative**  **Linear regression GWAS**  $\hat{b}_{zx}$**:** Allelic effect in phenotypic SD units of *x* | Estimated as $\hat{b}_{xy}$  Phenotypic SD units of *y* per phenotypic SD unit of *x* | $\hat{b}_{xy(logit \left( y \right):x)}$  ln(OR) for disease *y* per phenotypic SD of trait *x* |
| **Trait x is binary disease**  $\hat{b}_{zx(logit(x))}$**:** Allelic effect in ln(OR) of *x* | Estimated as $\hat{b}_{xy(y:logit(x))}$  $\hat{b}_{xy(y:liab(x))}=\frac{v_{K_{x}}}{K_{x}(1-K_{x})}\hat{b}_{xy(y:logit(x))}$  Phenotypic SD units of *y* per phenotypic liability SD unit of *x* | Estimated as $\hat{b}_{xy(logit\left( y \right):logit\left( x \right))}$  $\hat{b}_{xy(logit\left( y \right):liab(x))}=\frac{v_{K(x)}}{K_{x}(1-K_{x})}\hat{b}_{xy(logit\left( y \right):logit\left( x \right))}$  ln(OR) for disease *y* per phenotypic liability SD unit of *x*  $\hat{b}_{xy(liab\left( y \right):liab(x))}=\frac{v_{K_{x}}K_{y}(1-K_{y})}{v_{K_{y}}K_{x}(1-K_{x})}\hat{b}_{xy(logit\left( y \right):logit\left( x \right))}$  SD units of liability of *y* per SD unit of liability to *x* |

**3. Simulations to confirm derivations**

In order to confirm the derivations summarised in Supplementary Table 10, we performed a series of simulations. We simulated 100 SNPs from a binomial distribution *z* ~ B(2, *p*) with *p* being the allele frequency of a SNP *p* ~ U(0.01,0.5). We simulated an exposure phenotype under a liability-threshold model in (*n*=500,000 individuals) *x_liab_* = *b_zx_ z*+ *e*, *b_zx_* ~ N(0,1), $e\sim N(0,\sigma_{e}^{2})$, where $\sigma_{e}^{2}=\mathrm{var}(zb_{zx})(\frac{1}{R_{zx}^{2}}-1)$, with $R_{zx}^{2}$ was the variance in *x* explained by *z*. We set $R_{zx}^{2}=0.2$. *x* was then standardized with mean 0 and variance 1. Ten different sets of cases and controls were assigned to the underlying simulated liability according to various disease prevalences (*K*) from 0.01 to 0.1. The sample size of cases was $n_{cases}=nK$ and that of controls was $n_{controls}=n_{cases}(\frac{1}{P}-1)$, where *P* was sample prevalence, *P*~*U*(0.1, 0.5). The cases and controls were randomly sampled from the whole population. In the simulation, we estimated $\hat{b}_{zx(liab)}$ from the whole population by linear regression and $\hat{b}_{zx(logit)}$ from the ascertained sample by logistic regression. $\hat{b}_{zx(logit)}$ was then transformed to the liability scale. The simulation was replicated 1,000 times. The comparison of estimated and transformed $\hat{b}_{zx(liab)}$ is shown in the Supplementary Figure.

In the second set of simulations, a dichotomous exposure phenotype was simulated as above with 4 different prevalences (*K_x_* = 0.01, 0.04, 0.07 and 0.1) A outcome phenotype *y* with 200,000 individuals was simulated under the model $y_{liab}=x_{liab}b_{xy(liab)}+e_{y(liab)}$ _,_ where $e_{y\left( liab \right)}\sim N(0,\sigma_{e_{y(liab)}}^{2})$. $\sigma_{e_{y(liab)}}^{2}=\mathrm{var}(x_{liab}b_{xy\left( liab \right)})(\frac{1}{R_{xy}^{2}}-1)$ with $R_{xy}^{2}$ being the variance in *y* explained by *x*, $R_{xy}^{2}=0.05$. *y* was then standardized. We generated an ascertained sample for y using the same method as above. The four prevalences were the same as for disease *x*, *K_y_* = (0.01, 0.04, 0.07, 0.1). We estimated $\hat{b}_{xy(logit\left( y \right):logit(x))}$ by GSMR where $\hat{b}_{zx(logit(x))}$ and $\hat{b}_{zy(logit(y))}$ were estimated in the ascertained samples by logistic regression. $\hat{b}_{xy(logit)}$ was then transformed to $\hat{b}_{xy(liab)}$. The simulation was performed 1,000 times. Because both *x* and *y* were standardized, the expected value of $\hat{b}_{xy(liab)}=\sqrt{R_{xy}^{2}}=0.223$.

**Supplementary Table 10** shows the results from the simulation.

$\hat{b}_{xy(liab)}$ estimated from the logit scale. True $\hat{b}_{xy(liab)}$ is 0.223 which generated $\hat{b}_{xy(logit)}$ ranging from 0.156 to 0.304 depending of Kx, Ky combinations

| Disease prevalence | Scale | $K_{y}=0.01$ | $K_{y}=0.04$ | $K_{y}=0.07$ | $K_{y}=0.10$ |
| --- | --- | --- | --- | --- | --- |
| $K_{x}=0.01$ | logit | 0.214 (0.0007) | 0.180 (0.0004) | 0.165 (0.0003) | 0.156 (0.0002) |
|  | liability | 0.214 (0.0007) | 0.215 (0.0004) | 0.216 (0.0004) | 0.216 (0.0003) |
| $K_{x}=0.04$ | logit | 0.263 (0.0009) | 0.220 (0.0004) | 0.202 (0.0003) | 0.191 (0.0003) |
|  | liability | 0.219 (0.0007) | 0.220 (0.0004) | 0.220 (0.0004) | 0.220 (0.0003) |
| $K_{x}=0.07$ | logit | 0.287 (0.0009) | 0.240 (0.0005) | 0.221 (0.0004) | 0.209 (0.0003) |
|  | liability | 0.220 (0.0007) | 0.221 (0.0004) | 0.221 (0.0004) | 0.221 (0.0003) |
| $K_{x}=0.10$ | logit | 0.304 (0.0010) | 0.254 (0.0005) | 0.234 (0.0004) | 0.221 (0.0003) |
|  | liability | 0.220 (0.0007) | 0.221 (0.0004) | 0.221 (0.0004) | 0.221 (0.0003) |

The simulations show that the transformation to the liability scale from the logit scale is a good approximation of the true estimate of $\hat{b}_{xy(liab)}$. The estimate is slightly downward biased and the bias is greater when the prevalence of the exposure trait is low.

**Supplementary Figures**

Supplementary Figure 1. Results from simulations of

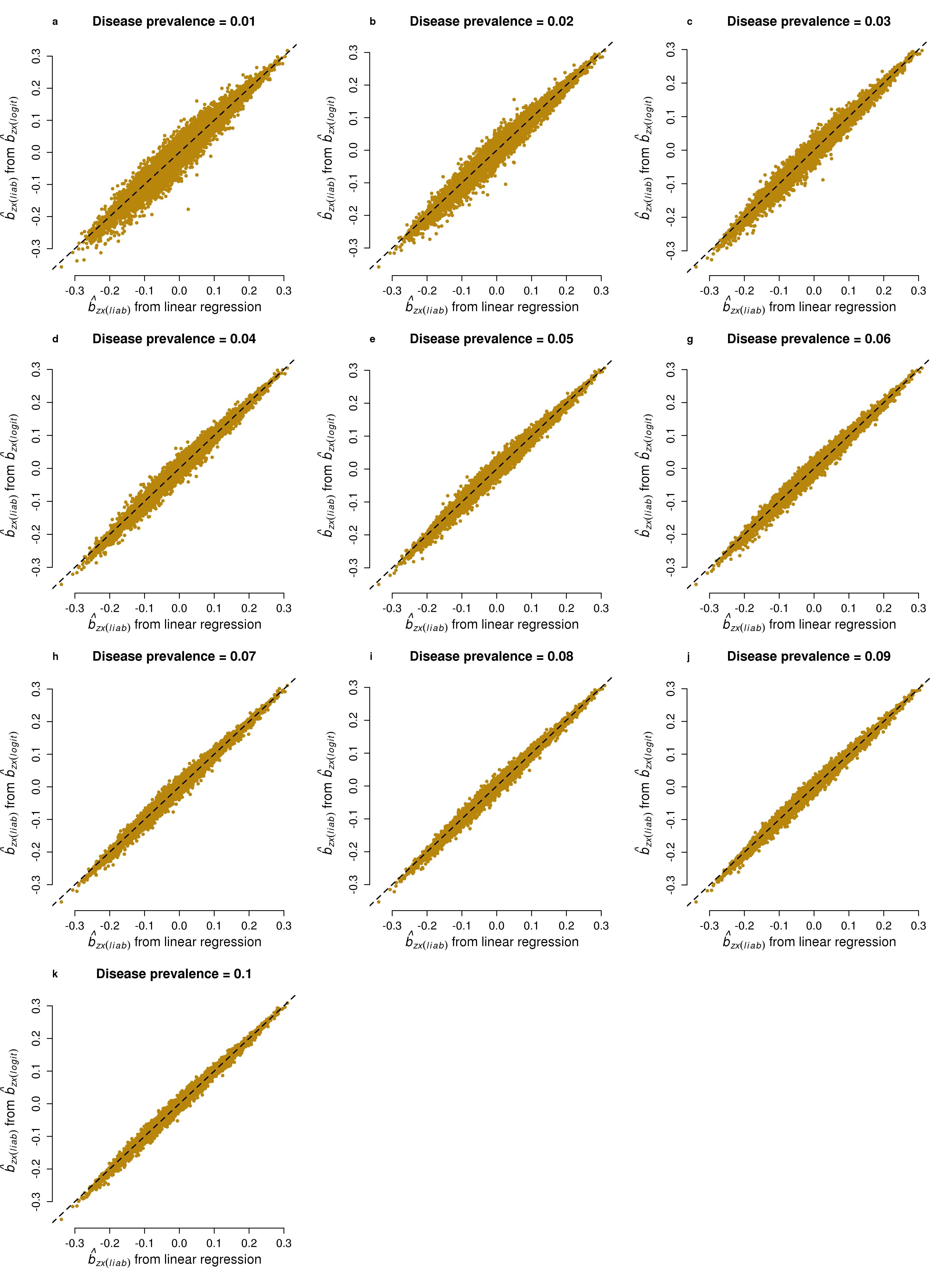

Supplementary Figure 2a-c. Significant SNPs from bipolar conditional analysis. All SNPs were not significant in the raw bipolar GWAS

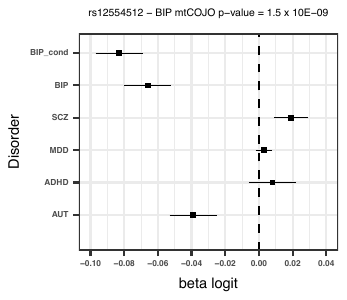

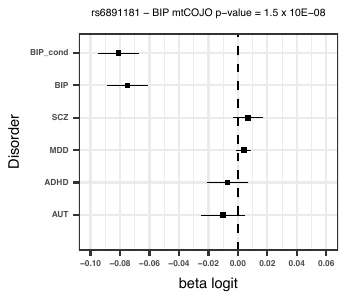

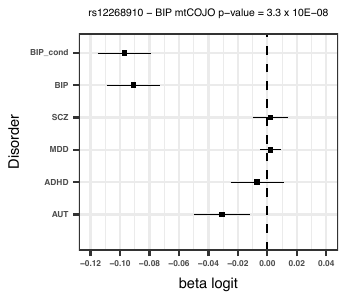

Supplementary Figure 3(a-e) Genome-wide significant SNPs from mtCOJO analysis of major depression with larger conditional effect sizes

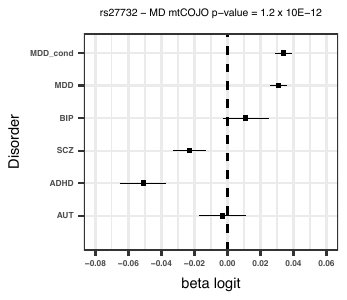

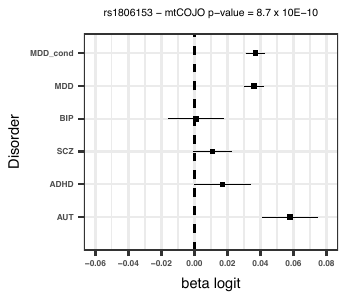

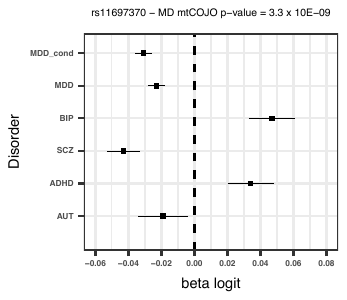

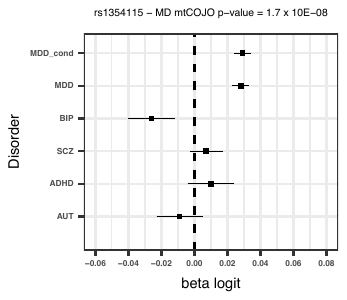

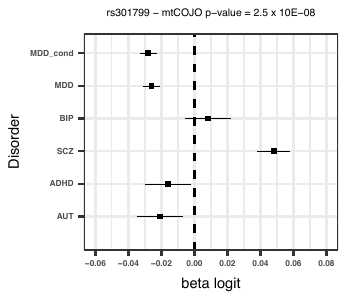

Supplementary Figure 4(a-e) Genome-wide significant SNPs from mtCOJO analysis of ADHD with larger conditional effect sizes.

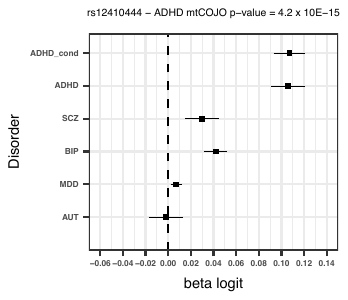

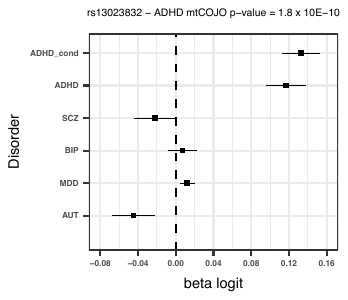

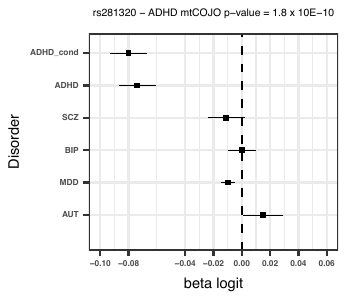

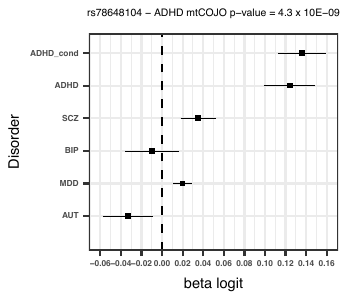

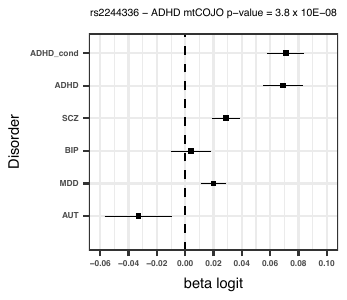

Supplementary Figure 5 Genome-wide significant SNP from mtCOJO analysis of AUT with larger conditional effect size.

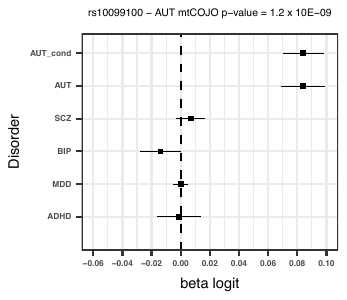

Supplementary Figure 6. Forest plot for mtCOJO analysis of rs3759384 – an eQTL for VPS29 – that was significant in SMR analysis.

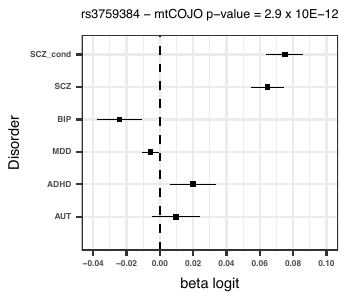

Supplementary Figure 7. Forest plot for mtCOJO analysis of rs2064853

– an eQTL for PCDHA7 in brain– that was significant in SMR analysis and has opposite effects on MD and SCZ

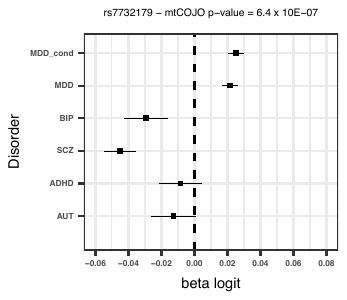

Supplementary Figure 8. Forest plot for mtCOJO analysis of rs2064853

– an mQTL for DNA methylation in the promotor of the CSE1L gene – that was significant in SMR analysis and has opposite effects on SCZ and BIP

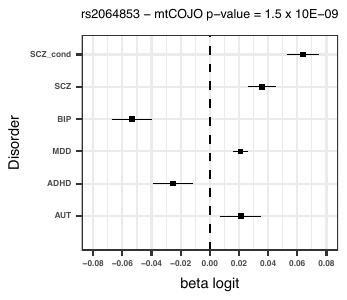

**Acknowledgments**

**ARIC**: The Atherosclerosis Risk in Communities Study is carried out as a collaborative study supported by National Heart, Lung, and Blood Institute contracts (HHSN268201100005C, HHSN268201100006C, HHSN268201100007C, HHSN268201100008C, HHSN268201100009C, HHSN268201100010C, HHSN268201100011C, and HHSN268201100012C), R01HL087641, R01HL59367 and R01HL086694; National Human Genome Research Institute contract U01HG004402; and National Institutes of Health contract HHSN268200625226C. The authors thank the staff and participants of the ARIC study for their important contributions. Infrastructure was partly supported by Grant Number UL1RR025005, a component of the National Institutes of Health and NIH Roadmap for Medical Research.

**CMC** (Synapse accession: syn2759792): CMC data were generated as part of the CommonMind Consortium supported by funding from Takeda Pharmaceuticals Company Limited, F. HoffmanLa Roche Ltd and NIH grants R01MH085542, R01MH093725, P50MH066392, P50MH080405, R01MH097276, RO1-MH-075916, P50M096891, P50MH084053S1, R37MH057881 and R37MH057881S1, HHSN271201300031C, AG02219, AG05138 and MH06692. Brain tissue for the study was obtained from the following brain bank collections: the Mount Sinai NIH Brain and Tissue Repository, the University of Pennsylvania Alzheimer’s Disease Core Center, the University of Pittsburgh NeuroBioBank and Brain and Tissue Repositories and the NIMH Human Brain Collection Core. CMC Leadership: Pamela Sklar, Joseph Buxbaum (Icahn School of Medicine at Mount Sinai), Bernie Devlin, David Lewis (University of Pittsburgh), Raquel Gur, Chang-Gyu Hahn (University of Pennsylvania), Keisuke Hirai, Hiroyoshi Toyoshiba (Takeda Pharmaceuticals Company Limited), Enrico Domenici, Laurent Essioux (F. Hoffman-La Roche Ltd), Lara Mangravite, Mette Peters (Sage Bionetworks), Thomas Lehner, Barbara Lipska (NIMH). GTEx (dbGaP accession: phs000424.v6.p1): The Genotype-Tissue Expression (GTEx) Project was supported by the Common Fund of the Office of the Director of the National Institutes of Health (commonfund.nih.gov/GTEx). Additional funds were provided by the NCI, NHGRI, NHLBI, NIDA, NIMH, and NINDS. Donors were enrolled at Biospecimen Source Sites funded by NCI Leidos Biomedical Research, Inc. subcontracts to the National Disease Research Interchange (10XS170), Roswell Park Cancer Institute (10XS171), and Science Care, Inc. (X10S172). The Laboratory, Data Analysis, and Coordinating Center (LDACC) was funded through a contract (HHSN268201000029C) to the The Broad Institute, Inc. Biorepository operations were funded through a Leidos Biomedical Research, Inc. subcontract to Van Andel Research Institute (10ST1035). Additional data repository and project management were provided by Leidos Biomedical Research, Inc.(HHSN261200800001E). The Brain Bank was supported supplements to University of Miami grant DA006227. Statistical Methods development grants were made to the University of Geneva (MH090941 & MH101814), the University of Chicago (MH090951, MH090937, MH101825, & MH101820), the University of North Carolina - Chapel Hill (MH090936), North Carolina State University (MH101819), Harvard University (MH090948), Stanford University (MH101782), Washington University (MH101810), and to the University of Pennsylvania (MH101822)

**GTEx** (dbGaP accession: phs000424.v6.p1): The Genotype-Tissue Expression (GTEx) Project was supported by the Common Fund of the Office of the Director of the National Institutes of Health (commonfund.nih.gov/GTEx). Additional funds were provided by the NCI, NHGRI, NHLBI, NIDA, NIMH, and NINDS. Donors were enrolled at Biospecimen Source Sites funded by NCI Leidos Biomedical Research, Inc. subcontracts to the National Disease Research Interchange (10XS170), Roswell Park Cancer Institute (10XS171), and Science Care, Inc. (X10S172). The Laboratory, Data Analysis, and Coordinating Center (LDACC) was funded through a contract (HHSN268201000029C) to the The Broad Institute, Inc. Biorepository operations were funded through a Leidos Biomedical Research, Inc. subcontract to Van Andel Research Institute (10ST1035). Additional data repository and project management were provided by Leidos Biomedical Research, Inc.(HHSN261200800001E). The Brain Bank was supported supplements to University of Miami grant DA006227. Statistical Methods development grants were made to the University of Geneva (MH090941 & MH101814), the University of Chicago (MH090951, MH090937, MH101825, & MH101820), the University of North Carolina - Chapel Hill (MH090936), North Carolina State University (MH101819), Harvard University (MH090948), Stanford University (MH101782), Washington University (MH101810), and to the University of Pennsylvania (MH101822)
